# Virome responses to heating of a forest soil suggest that most dsDNA viral particles do not persist at 90°C

**DOI:** 10.1101/2024.05.01.592093

**Authors:** Sara E. Geonczy, Luke S. Hillary, Christian Santos-Medellín, Jane D. Fudyma, Jess W. Sorensen, Joanne B. Emerson

## Abstract

Many fundamental characteristics of soil viruses remain underexplored, including the effects of high temperatures on viruses and their hosts, as would be encountered under disturbances like wildland fire, prescribed burning, and soil solarization. In this study, we leveraged three data types (DNase-treated viromes, non-DNase-treated viromes, and 16S rRNA gene amplicon sequencing) to measure the responses of soil viral and prokaryotic communities to heating to 30°C, 60°C, or 90°C, in comparison to field and control conditions. We investigated (1) the response of dsDNA viral communities to heating of soils from two horizons (O and A) from the same forest soil profile, (2) the extent to which specific viral taxa could be identified as heat-sensitive or heat-tolerant across replicates and soil horizons, and (3) prokaryotic and virus-host dynamics in response to heating. We found that both viral and prokaryotic communities responded similarly to the treatment variables. Community composition differed most significantly by soil source (O or A horizon). Within both soil horizons, viral and prokaryotic communities clustered into three groups, based on beta-diversity patterns: the ambient community (field, control, and 30°C samples) and the 60°C and 90°C communities. As DNase-treated viromic DNA yields were below detection limits at 90°C, we infer that most viral capsids were compromised after the 90°C treatment, indicating a maximum temperature threshold between 60°C and 90°C for most viral particles in these soils. We also identified groups of heat-tolerant and heat-sensitive vOTUs across both soil sources. Overall, we found that over 70% of viral populations, like their prokaryotic counterparts, could withstand temperatures as high as 60°C, with shifts in relative abundance explaining most community compositional differences across heating treatments.

## INTRODUCTION

Soil viral ecology is a growing field that explores how viruses interact with their hosts and the environment, which can help characterize their role in soil functions (Emerson, 2019; Kimura et al., 2008; Kuzyakov and Mason-Jones, 2018; Trubl et al., 2020). Many soil functions, particularly biogeochemical cycling, are driven by microbial processes that immobilize, mineralize, and otherwise transform key nutrients (Kuypers et al., 2018; Lladó et al., 2017), with viral infection presumed to impact these processes in largely unknown ways. Prokaryotic viruses target bacteria and archaea, impacting host metabolic activity and leading to cell lysis through various forms of infection dynamics (Labrie et al., 2010). Viruses are also theorized to play a direct part in nutrient cycling, separate from their impact on microorganisms, for example as stoichiometrically enriched sources of phosphorus (Jover et al., 2014). Many basic characterizations of soil viruses remain ripe for further exploration, including the impact of temperature on viral ecology.

The temperature range that an organism can tolerate while remaining viable is a defining characteristic that influences ecological responses to changing environmental conditions and can impact biogeographical patterns. For soil organisms, certain disturbances can increase soil temperatures outside the range of an average diurnal cycle. Soil solarization can increase temperatures up to more than 50°C at 5 cm depth (Oz et al., 2017), soils under forest slash burn piles can reach up to 138 ± 28°C at 5 cm depth (Busse et al., 2013), and controlled surface burns can increase soil temperatures to over 400°C at 2 cm (Massman and Frank, 2004). Temperature tolerance determines which organisms can or cannot survive these disturbances and drives community compositional changes (Siliakus et al., 2017). While more is known about prokaryotic temperature tolerances in soil (Donhauser et al., 2020; Pingree and Kobziar, 2019), studying the temperature tolerances of the viruses of these prokaryotes can help us to better understand virus-host dynamics and post-disturbance ecological responses.

High temperatures are known to inactivate viruses and compromise viral capsids, releasing nucleic acids into the environment (Bertrand et al., 2012; Jończyk et al., 2011; Kimura et al., 2008; Williamson et al., 2017). Upper thresholds for persistence vary, and prokaryotic viruses have been found in extreme thermal environments, serving as the only entities that can continue to apply biological pressure on microorganisms at high temperatures (Breitbart et al., 2004). Studies of sewage and sludge have also shown that phages can be more resistant to thermally-induced inactivation than bacteria (Mocé-Llivina et al., 2003). In dairy production studies, temperate phages of *Lactococcus* spp. have been shown to survive temperatures as high as 97°C after 5 minutes of heating, with even more heat-resistant dairy bacteriophages still being discovered (Atamer et al., 2009; Buzrul et al., 2007; Guglielmotti et al., 2012). However, little is known about prokaryotic viruses in soil and their responses to soil heating.

The degree of temperature change from heating in soil depends on a suite of physicochemical characteristics, including soil moisture, organic matter content, and mineralogy (Bruns et al., 2020; Dooley and Treseder, 2012; Janzen and Tobin-Janzen, 2008; Lucas-Borja et al., 2019; Mataix-Solera et al., 2009). Soil also does not heat evenly, and there is evidence of stratified microbial survival in soil based on depth (Palmer et al., 2023). Studies in soils show that there is a range of prokaryotic survival thresholds that varies by temperature and duration of heating, likely owing to different adaptations of prokaryotes, as well as the heterogeneity of soils which impacts the extent of heat penetration (Pingree and Kobziar, 2019). For instance, organic soils that typically comprise the upper section of a soil profile, considered the O horizon, have high organic matter content. Mineral soils, which have lower organic matter, are more dominated by the mineral fraction of soil. In forests, mineral soils tend to underlie an O horizon, typically considered the A horizon (Neary et al., 2005; Osman, 2013). This makes it important to study the impact of increased temperature on various soil types, with special attention to different horizons within a soil profile that have distinct properties and would intrinsically respond differently to heating (Busse et al., 2014; Certini, 2005; Johnson and Turner, 2019; van Wagtendonk et al., 2018).

Fortunately, the ability to investigate the temperature tolerance of viruses in soil is enabled via viromics. Viromics is a size-fractionated metagenomic approach that enriches for virus-like-particles (VLPs), in many cases specifically enriching for dsDNA viruses (Santos-Medellin et al., 2021; Trubl et al., 2020). Viral particles are brought into solution via mechanical and chemical means to remove most free virions from the soil matrix, including those that were adhered to soil particles or occluded in aggregate pores. DNase treatment, the addition of a DNA-degrading enzyme, cleans samples by removing the vast majority of DNA occurring outside a virion capsid prior to DNA extraction (Göller et al., 2020; Sorensen et al., 2021). This leaves samples that are enriched in extracellular virions (not prophages or viruses in the intracellular stage of their replication cycle) with intact capsids that are theoretically viable, or at least more likely to be viable than virions with compromised capsids. Thus, DNase-treated viromics is a way to measure active viral community dynamics (Sorensen et al., 2021). However, when considering viromes that have not been treated with DNase, sequences can be recovered not only from virions that remained intact and potentially viable, but also from viruses that have been compromised (Santos-Medellín et al., 2023), which can provide further insights into persistence under changing environmental conditions.

Viromes, whether treated with DNase or not, also enrich for small prokaryotic cells that are able to pass through a 0.22 µm filtration step, including those belonging to the superphylum Patescibacteria, which are ultra-small bacteria with small genome sizes (Nicolas et al., 2021; Santos-Medellín et al., 2023; Tian et al., 2020). Thus, 16S rRNA gene amplicon sequencing of viromes can also reveal dynamics of small bacteria, in which the signal is otherwise obscured with total DNA extraction. Viromes not treated with DNase can also potentially reveal microbial mortality through characterization of relic DNA of prokaryotes (Santos-Medellín et al., 2023). This can be especially useful if paired with sequencing results from total DNA, which would include prokaryotes both potentially active (cells intact) and dead.

In this study, we leveraged these three sample types (DNase-treated viromes, non-DNase-treated viromes, and total DNA) to measure the responses of soil viral and prokaryotic communities to heating treatments of target temperatures between 30°C and 90°C and to deduce virus-host dynamics. Primarily, we used DNase-treated viromes to characterize presumed active viral communities, and we used total DNA for prokaryotic communities. We used non-DNase-treated viromes to yield additional insights into the inactive/relic fraction of viruses and prokaryotes (e.g., for additional evidence as to which species succumbed at which temperature thresholds), as well as to infer the responses of small bacteria to increasing temperature. From both total DNA and the non-DNase-treated viromes, we generated 16S rRNA gene amplicon sequencing data to characterize prokaryotic communities. The three main questions that we address in this study are: (1) How do dsDNA viral communities respond to heating of soils from two horizons (O and A) from the same soil profile? (2) To what extent are specific viral taxa consistently heat-sensitive or heat-tolerant across replicates and soil types? and (3) What are the prokaryotic and virus-host dynamics in response to heating?

## METHODS

### Soil Collection

Soil for this study was collected from Blodgett Forest Research Station in California, USA (38°54.905’ N, 120°39.640’ W, elevation 4299 ft) on April 18, 2022. This location was chosen due to an ongoing field experiment in the same area set up one year prior, which will study the impact of prescribed burning on soil viral ecology. Building on evidence that spatial variability obscures treatment effects in soil virus studies (Durham et al., 2022; Fudyma et al., 2024; Santos-Medellín et al., 2022), we sought to constrain spatial variability as much as possible, collecting two soil types (O and A horizon) from one location. The location was picked because of its soil profile characteristics, where the O horizon corresponded to a 0-3 cm depth of the soil profile with distinct separation along the 3 cm depth between the O horizon and underlying A horizon. We determined the soil profile using a soil probe. To collect 2 kg of each soil (the amount we had calculated to satisfy all treatment and metadata requirements), the total excavation site approximated a 40 x 70 x 6 cm block, with O horizon soil from the top 3 cm and A horizon soil from the bottom 3 cm. We homogenized and sieved (8 mm) each soil type.

### Selection of Experimental Parameters

For experimental soil heating, we chose 10°C as the control temperature, which approximated field temperatures without a heating disturbance. To determine this temperature, temperature measurements were taken just prior to soil collection via a soil thermometer at 3 and 6 cm depth in random locations within the soil collection area (Supplementary Table 2). In addition, temperatures were measured at six nearby plots (four burn plots and two control plots) over a period of 6 days before and after a prescribed burn in the previous year as part of the aforementioned ongoing study. For this, we used temperature probes (Extech SDL200 4-Channel Data Logging Thermometer, Extech TP873 Bead Wire Type K Temperature Probe, -30 to 300°C) inserted at 1, 2, 4, and 6 cm in soil profile with at least one logging unit per plot (Supplementary Table 3), allowing us to select three biologically and ecologically relevant temperatures for the heating treatments. We chose 90°C for the highest temperature, due to evidence of some viral persistence at these temperatures from the literature (discussed in the Introduction), 60°C for the second-highest temperature, as prokaryotic community shifts would be expected due to some bacterial mortality at that temperature (Busse et al., 2014), and 30°C for the lower temperature, as this represents elevated temperatures seen in some heating disturbances (solarization, surface burning) but is lower than most known survival thresholds for prokaryotes and virions. A period of 30 minutes was selected for the heating duration to sufficiently heat up the soil to a stabilized temperature and to mimic shorter-term heating disturbances that would happen naturally in prescribed burning.

### Laboratory microcosms subjected to soil heating

The experimental treatments were as follows: six samples (three replicates per horizon) were processed for DNA extraction the day after field collection without any heating treatment, after storage at 4°C for approximately 12 hours. These are considered “field” samples because they were processed shortly after field sampling. Each of the other sample sets (three samples per horizon per set) were randomly divided into two 36-hour treatment periods that occurred in succession. This time was chosen so that samples could be processed within a two-day period (12 samples per day). At the beginning of each treatment period, samples were placed in a light-penetrable incubator (New Brunswick Excella E-24R) at 10°C to maintain soil at the control temperature. At the beginning of the 36 hours, samples were subjected to 30 minutes of heating by removing tubes from the incubator and placing them in heat blocks (VWR 13259-254) that were pre-heated to one of three temperatures (30, 60, or 90°C).

Soil samples were added up to the 20 mL line of 50 mL bioreaction centrifuge tubes (VWR # 76211-286), which maximized the amount of soil exposed to the heating block. Due to differences in bulk density (0.15 g/cm^3^ for the O horizon soil and 0.39 g/cm^3^ for the A horizon soil), each sample weighed on average 6.39 g for the organic soil and 9.77 g for the mineral soil. The tubes had vented caps with 0.22 µm membranes, which allowed for free gas exchange while maintaining aseptic conditions. After 30 minutes, the samples were returned to the 10°C incubator. A sample set remained in the 10°C incubator for the entire 36 hours and is considered the “control” sample set. For each treatment, an additional sample (not processed for downstream measurements) was used for measuring the soil temperature in the center of the tube at the start and at 10 min intervals throughout the 30-minute heating incubation. Once the 36 hours were complete, the samples were collected for DNA extraction. Samples were heated and processed in a random order to minimize batch effects.

### Viromic DNA extraction and quantification

We extracted viromic DNA from all samples, as previously described (Santos-Medellín et al., 2022). Briefly, to each soil sample, we initially added 9 mL of protein-supplemented phosphate-buffered saline solution (PPBS: 2% bovine serum albumin, 10% phosphate-buffered saline, 1% potassium citrate, and 150 mM MgSO4, pH 6.5). Samples were vortexed until homogenized and then placed on a horizontal shaker at 300 rpm at 4°C for 10 min. Supernatant was transferred into a new tube and then 9 mL of PPBS was added to the soil remaining in the original tube, repeating the process for a total of three soil resuspensions and elutions, ending with a total of three volumes of supernatant pooled together in a new centrifuge tube (approximately 27 mL). The pooled supernatant was then centrifuged at 8 min at 10,000 x g, 4°C, with the progressively cleaner supernatant collected and the concentrated soil particles discarded, for a total of three centrifugation and supernatant collection steps. The final supernatant was filtered through a 5 µm polyethersulfone (PES) membrane filter (brand: PALL) to filter out larger soil particles and then the effluent went through a 0.22 µm PES membrane filter (brand: PALL) to exclude most cellular organisms. Virus-sized particles and DNA remaining in the effluent were concentrated into a pellet using the Optima LE-80K ultracentrifuge with a 50.2 Ti rotor (Beckman-Coulter) at 32,000 x g at 4°C for 2 h 25 min. The pellet was resuspended in 200 µL of 0.02 µm filtered ultrapure water and split into two tubes, with one half treated with DNase (briefly, incubation at 37°C for 30 min after adding 10 units of RQ1 RNase-free DNase, Promega) and one half not treated with DNase (considered non-DNase-treated virome samples). For all virome samples, we extracted DNA using the DNeasy PowerSoil Pro kit (Qiagen), following the manufacturer’s protocol, with the addition of a 10 min incubation at 65°C prior to cell lysis, which also served as the ‘stop’ incubation for the DNase treatment. DNA quantification was performed using the Qubit dsDNA HS Assay and Qubit 4 fluorometer (Thermo Fisher), and samples with yields above 0.1 ng/µl were prepared for library construction and sequencing.

### Total DNA extraction

For each sample, 0.25 g of soil was added directly to the DNeasy PowerSoil Pro kit (Qiagen) for DNA extraction, following the manufacturer’s protocol, with the addition of a 10 min incubation at 65°C prior to cell lysis.

### Shotgun virome library preparation and sequencing

Sequencing of all viromes was performed by the DNA Technologies and Expression Analysis Core at the University of California, Davis Genome Center. Metagenomic libraries were constructed with the DNA Hyper Prep kit (Kapa Biosystems-Roche) and sequenced (paired-end, 150 bp) using the Illumina NovaSeq S4 platform to a requested depth of 10 Gbp per virome.

### Amplicon library preparation and sequencing

We performed 16S rRNA gene amplicon sequencing on all total DNA extractions and on all the non-DNase treated viromes. We followed Earth Microbiome Project’s PCR protocol (Thompson et al., 2017), which included an initial denaturation step at 94°C for 3 min, 35 cycles of 94°C for 45 s, 50°C for 60 s, and 72°C for 90 s, and a final extension step at 72°C for 10 min. We cleaned libraries with AmpureXP magnetic beads (Beckman Coulter), quantified amplicons with a Qubit 4 fluorometer (Thermo Fisher), and then pooled all bead-purified products together in equimolar concentrations. Libraries were sequenced by the DNA Technologies and Expression Analysis Core at the University of California, Davis Genome Center, paired-end (250 bp) on the Illumina MiSeq platform.

### Soil physicochemical measurements

We calculated gravimetric soil moisture nine times for each soil horizon, by weighing out 10 g of soil into aluminum weigh dishes and measuring the weight after 24 hours in a drying oven at 105°C. Three replicate samples of field-condition soil (not dried) of each horizon and one sample for each post-heat treatment (at 30, 60, and 90°C temperatures) were sent to Ward Laboratories (Kearney, Nebraska, USA) for a suite of soil chemistry analyses under the Haney Soil Health Analysis package (“Haney Test Interpretation Guide v1.0,” n.d.), which is summarized in Supplementary Table 4. Virome bioinformatic processing:

Raw reads were trimmed using Trimmomatic v0.39 (Bolger et al., 2014) with a minimum q-score of 30 and a minimum read length of 50 bases. We used BBMap v39.01 (Bushnell, 2014) to remove PhiX sequences. Each virome was assembled separately with MEGAHIT v1.2.9 (Li et al., 2015) in the metalarge mode with a minimum contig length of 10,000 bp. Assembled contigs were analyzed with VIBRANT v1.2.0 (Kieft et al., 2020) with the -virome flag to identify viral contigs. All output viral contigs (low, medium, and high quality) from all samples were iteratively clustered to form a representative, dereplicated set of vOTUs. We first used CD-HIT v4.8.1 (Huang et al., 2010) to cluster viral contigs at 95% shared nucleotide identity and 85% alignment coverage (breadth). We then consolidated the resulting contigs into species-level viral operational taxonomic units (vOTUs) with dRep v3.2.2 (Olm et al., 2017), using the default algorithm for secondary clustering comparisons, default ANI threshold to form secondary clusters (0.95), and 0.85 minimum level of overlap between genomes when doing secondary comparisons. Quality-filtered raw reads from all viromes were then mapped against the dereplicated set of vOTUs using Bowtie 2 v2.4.1 (Langmead et al., 2009) in sensitive mode. Lastly, we used CoverM v0.5.0 (Woodcroft, 2023) to quantify vOTU abundance and generated a trimmed mean coverage table and a count table for downstream analyses (0.75 minimum covered fraction). Host taxonomy of the entire set of dereplicated vOTUs was predicted using iPHoP v1.1.0 (Roux et al., 2023), with default parameters (minimum cutoff score of 90).

### 16S rRNA gene amplicon sequence bioinformatic processing

We used DADA2 v1.12.1 (Callahan et al., 2016) to demultiplex 16S rRNA gene amplicon reads and perform quality assessment, read merging, and chimera removal. Amplicon sequence variants (ASVs) were assigned taxonomy by using the DADA2 RDP classifier, using the SILVA database v138 as reference (Quast et al., 2013).

### Statistical and ecological analyses

We used R v4.3.0 to perform all ecological and statistical analyses (R Core Team, 2023). Input for all analyses of viral communities or vOTUs was the trimmed mean coverage vOTU table. We normalized the vOTU abundance table by using the decostand function (method = “total”, MARGIN = 2) from the Vegan package v2.6-4 (Oksanen et al., 2022) and removed singletons (vOTUs detected in only one sample). We used a DADA2-generated abundance table for 16S rRNA gene amplified single variants (ASVs), filtered out mitochondria and chloroplasts, and removed singletons (ASVs that only appeared in a single sample). We used the vegdist function (method = “bray”) to calculate Bray-Curtis dissimilarities and the adonis function to perform PERMANOVAs, with both functions from Vegan. We used the cmdscale function to perform multidimensional scaling and retrieve true eigenvalues and calculate PCoA points. We used ComplexUpset v1.3.5 to create upset plots using species abundance tables that were transformed into presence-absence tables (Krassowski, 2021). To determine enrichment of vOTUs along the temperature gradient, we performed a differential abundance analysis, using the tool DESeq2 v1.38.3 (Love et al., 2014). We performed differential abundance comparisons among all temperatures within horizon and sample type subgroups, then adjusted P-values using the Bonferroni method to determine overall significance of differentially abundant contigs. To identify trend groups among these contigs, we used k-means clustering.

### Gene-sharing Network

We predicted protein content for each vOTU using Prodigal v2.6.3 (metagenome mode) (Hyatt et al., 2010) and used the amino acid file output for gene-sharing network analysis with the tool vConTACT v2.0 (Jang et al., 2019). We used Diamond for protein alignment (Buchfink et al., 2021) and the MCL algorithm for clustering proteins. We used the Fruchterman-Reingold algorithm from the Ggally package (Schloerke et al., 2024) to visualize the resulting network, additionally highlighting a subnetwork with neighborhoods that were enriched in significantly differentially abundant vOTUs. All scripts and intermediate files are available at https://github.com/seugeo/soilheating.

## RESULTS

### Study Design and Dataset Features

To evaluate the extent to which extractable viral and prokaryotic community composition differed in response to heating treatments and to constrain the threshold temperature(s) for likely decay of most viral particles, we performed a laboratory heating experiment on soils from two horizons (O and A horizons) from a mixed conifer forest (Figure 1A). The soil from each horizon was divided into five sample groups, each with three technical replicates (Figure 1B). The ‘field’ sample group was processed within 24 hours of sample collection for viromics and total DNA extraction to capture viral and prokaryotic communities present in the field. The remaining sample groups were subjected to a 36-hour incubation at 10°C, with three of the sample groups elevated to higher temperatures (30, 60, or 90°C), for 30 minutes at the beginning of the incubation. The fourth set of samples remained in the 10°C incubation for the entire 36 hours and is considered the control group. The temperature at the center of the test tube reached 84.4°C for the O horizon soil and 83.0°C for the A horizon soil when placed in a 90°C heating block, 58.0°C for the O horizon soil and 57.9°C for the A horizon soil in the 60°C heating block, and 29°C for the O horizon soil and 28.8°C for the A horizon soil in the 30°C heating block (Figure 1C). As expected, the soil did not heat evenly, so when reporting the intended temperature as the heating block temperature (e.g. 90°C), we acknowledge that not all of the soil reached that temperature.

**Figure 1.**
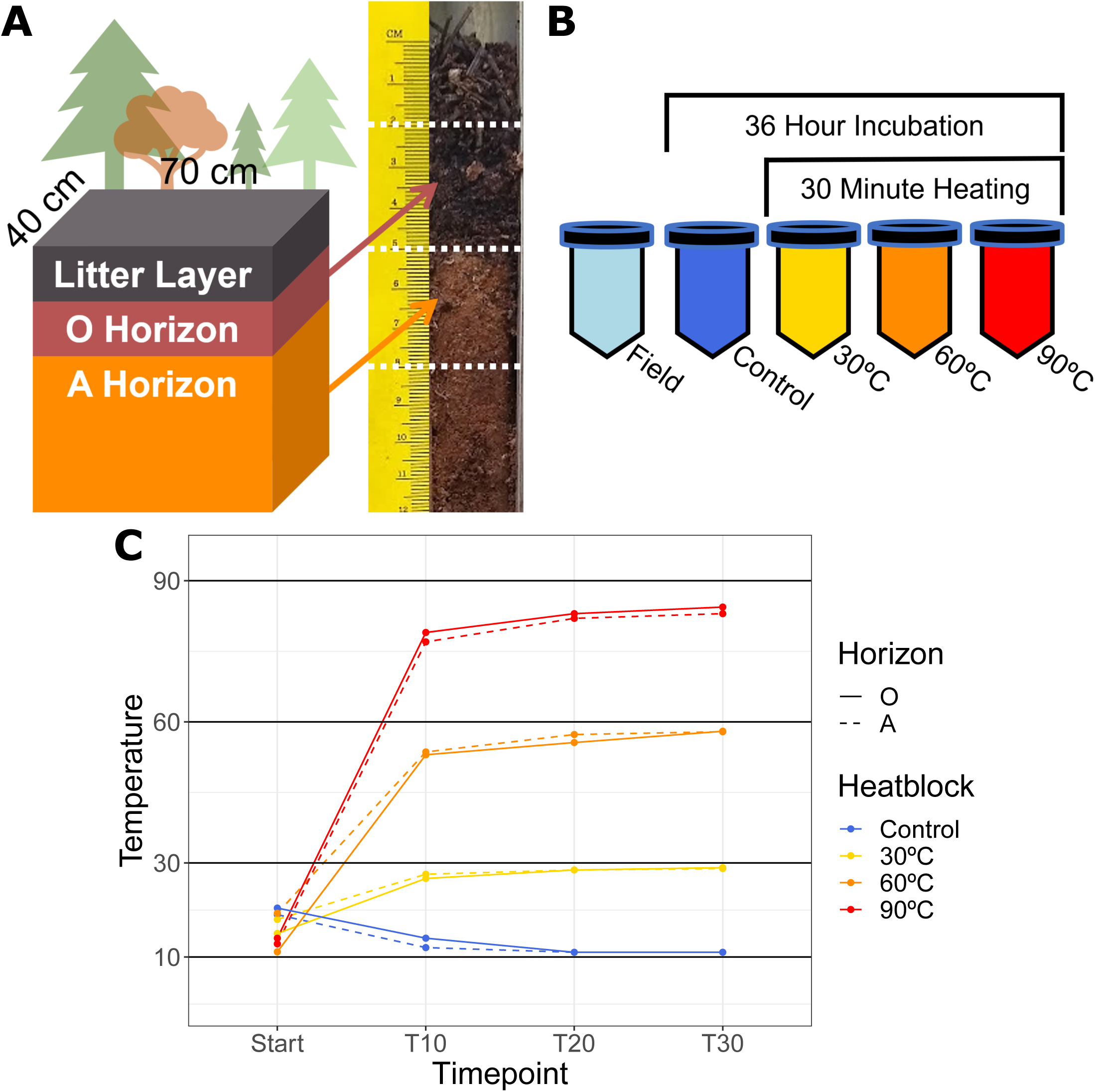
Dataset Features. (**A**) Sample collection source: graphic depicts surface area of soil collection, arrows point to photo of a representative soil probe in the sample collection area, showing soil quality and sample depth of each horizon. The color of the horizons in the graphic is maintained in subsequent figures. (B) Experimental design: each of five treatments is represented by a unique color that will be maintained throughout subsequent figures. Field samples were processed without an incubation step, while all other samples were incubated for 36 hours, with three sample sets elevated to different target temperatures (30, 60, or 90°C) over 30 minutes during the 36-hour incubation. (C) Microcosm soil temperature over time (10-minute intervals), from start to 30 minutes (T30) for all incubated samples. Control samples were not added to a heat block, and the other samples were placed in a heat block that was preheated to the indicated temperature.

We sought to characterize the temperature responses of viral particles, microorganisms, and relic DNA to determine which viruses and organisms likely remained viable and which likely did not persist across the temperature gradient. We performed a suite of DNA extraction and sequencing approaches to compare the viral particle, prokaryotic, and relic DNA fractions. Two types of viromes were prepared by sequencing DNA from the post-0.22 µm (viromic) DNA fraction: prior to DNA extraction, each <0.22 µm viral concentrate was split in half, with one half treated with DNase (‘DNase-treated’), and the other half not treated with DNase (henceforth referred to as ‘non-DNase treated’), for a total of 60 extracted viromes from 30 samples. Based on our prior work (Santos-Medellín et al., 2023; Sorensen et al., 2021), the DNase-treated viromes capture primarily intact viral particles, whereas the non-DNase treated viromes capture a combination of intact viral particles, compromised viral particles, extracellular DNA potentially reflecting dead microbes and viruses, and small cells that passed through the 0.22 µm filter. DNA yields were below detection limits (less than 0.1 ng/µl) from all six DNase-treated viromes (both soil horizons) at 90°C, and thus we were able to sequence 54 of the 60 prepared viromes (Figure 3G). From the 54 sequenced viromes, 145,914 contigs >10 kbp were assembled, 141,638 of which were identified as viral via VIBRANT (Kieft et al., 2020). These viral contigs clustered into 18,869 representative vOTU sequences, and we analyzed a final dataset of 18,538 vOTUs after removing singletons (vOTUs only appearing in a single sample).

To investigate prokaryotic communities, we generated 16S rRNA gene amplicon sequences from both total DNA, which captures a combination of active, dormant, and dead prokaryotes (Burkert et al., 2019; Carini et al., 2016), and from non-DNase viromes, which would include relic (extracellular) DNA that co-precipitated with viral particles, as well as an enrichment of small-celled microorganisms (Santos-Medellín et al., 2023, 2022). We were able to perform 16S rRNA gene amplicon sequencing on all 30 total DNA samples and all 30 non-DNase viromes, together yielding 24,270 ASVs, which reduced to 6,679 ASVs after rarefying to a common depth of 25,000 reads per sample and removing singletons.

### Viral and prokaryotic community composition differed most significantly by soil source (between O and A horizons) and secondarily along a temperature gradient

Considering the full dataset of 54 sequenced viromes, soil viral communities differed most significantly by soil source (between O and A horizons, P < 0.001 by PERMANOVA) (Figure 2A). When considering DNase-treated viromes, the largest fraction of vOTUs was shared between the two horizons (39%), but the fraction that was distinct to each horizon was still substantial (32% and 29% in O and A horizons, respectively) (Figure 2B), showing compositional distinction between the two soil sources. Despite close physical proximity, the distinct physicochemical features of each soil horizon (*e.g.*, microbially active carbon, pH, organic matter, available nitrogen) presumably contributed to the differences in viral community composition (Figure 2C). A robust comparison of temperature responses by soil horizon would have required biological replicates from different locations, which was not our goal in this study. Rather, our interest was to compare two distinct soil sources, empirically confirmed here as distinct in both viral and physicochemical composition, to determine whether two different soil sources would yield similar measured responses to temperature.

**Figure 2.**
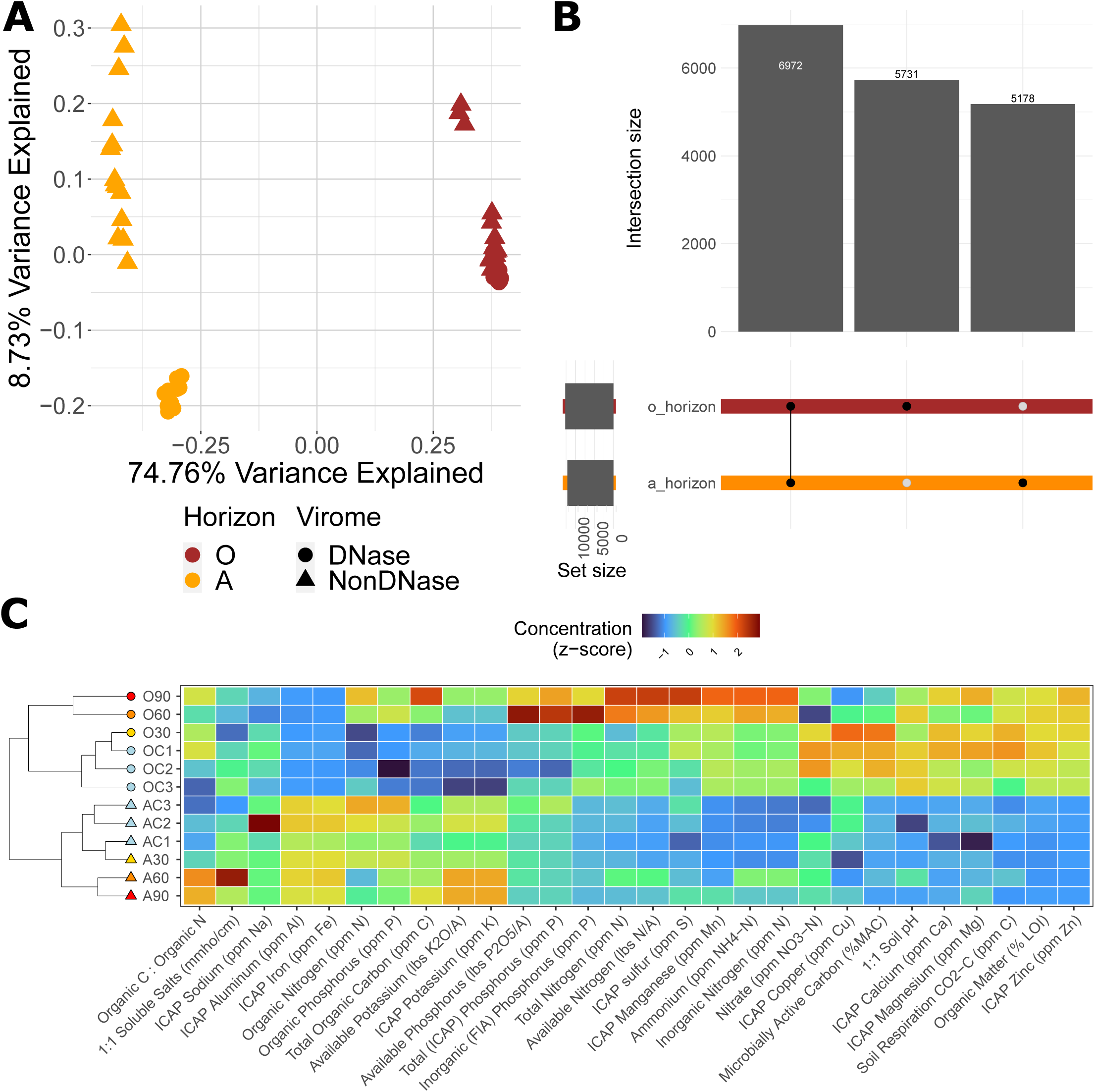
Differences in viral community composition by soil type. (**A**) Unconstrained analyses of principal coordinates (PCoA) performed on vOTU Bray-Curtis dissimilarities calculated across all viromes, with colors indicating soil source and shapes indicating virome type. (**B**) UpSet plot of vOTU detection in each horizon in DNase-treated viromes, with filled circles indicating detection in a particular horizon and intersection size indicating the number of vOTUs with a given detection pattern (one of the two horizons or both). (C) Hierarchical clustering of soil samples based on soil chemical data, with three replicates of each horizon in the field treatment and one replicate of each horizon in the 30, 60, and 90°C treatments. Sample labels begin with horizon (A or O), followed by “F” for field or the incubation temperature. The corresponding heatmap shows the z-transformation of each environmental measurement across samples.

Within each soil source (O or A horizon), viral communities differed significantly based on temperature treatment (P < 0.001 for O horizon, P < 0.001 for A horizon by PERMANOVA) (Figures 3A&B). In both horizons, viromes significantly separated into three treatment groups: one group with field, control, and 30°C viromes (hereafter referred to as the “ambient” community), a second with 60°C viromes, and a third with 90°C viromes, noting that the 90°C temperature group only included non-DNase viromes in both horizons, as DNase-treated viromes yielded no detectable DNA at that temperature. In DNase-treated viromes from both horizons, the vast majority of vOTUs was detected in all treatment groups (82% in O horizon and 73% in A horizon), suggesting that at least some members of most viral populations persisted up to 60°C (the highest temperature from which DNase-treated viromes were recovered). The next largest group of vOTUs (8% in O horizon and 7% in A horizon) was present in all treatments, except for the highest temperature, which in this case was 60°C (Figures 3C&D and Supplementary Figures 1A-D), suggesting temperature-induced decay of those viral populations between 30°C and 60°C. These results show that in the DNase-treated viromes, the presence or absence of unique vOTUs is likely not the main driver separating ambient and 60°C communities.

**Figure 3.**
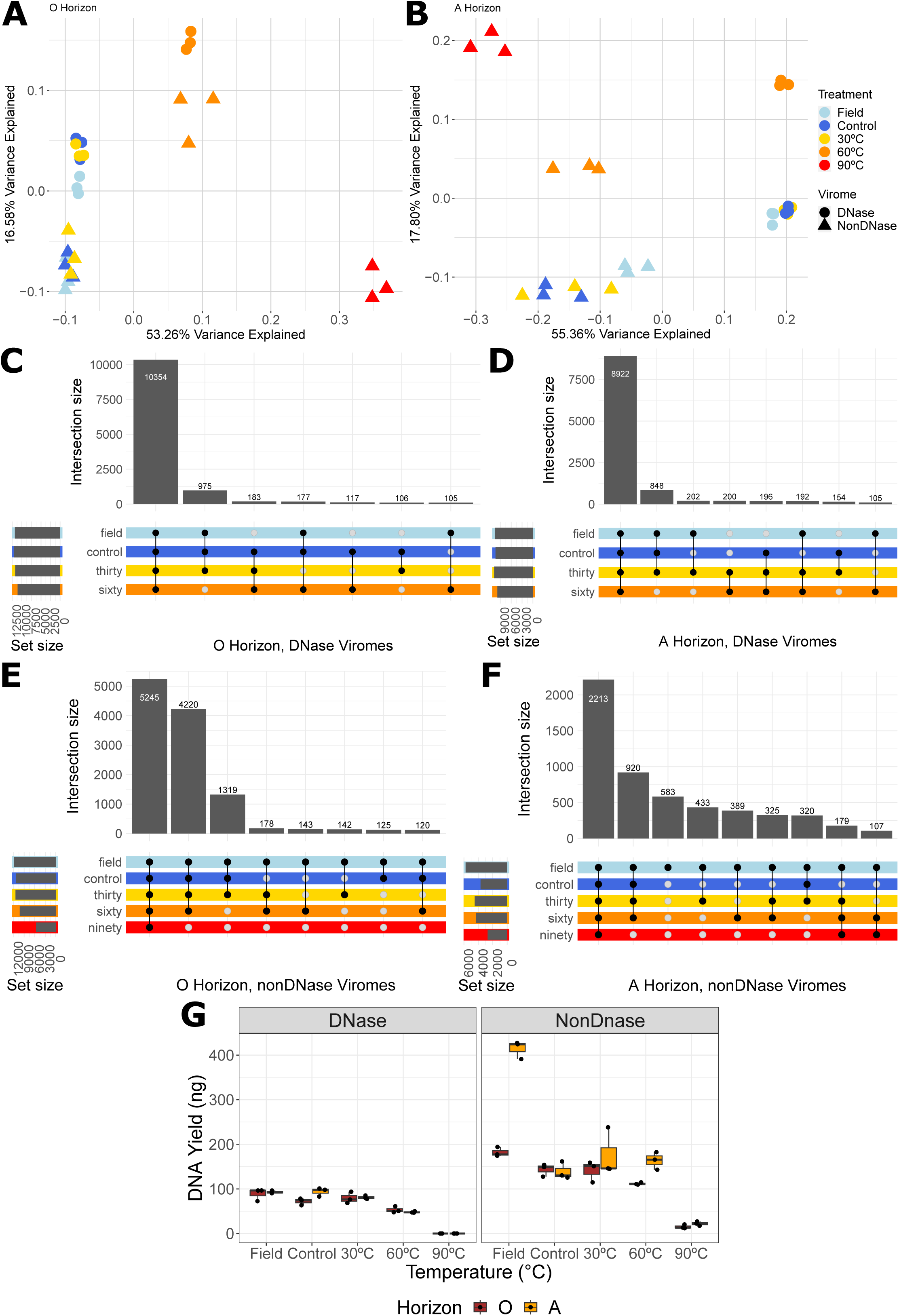
Viral community composition shifts based on heating treatments. Unconstrained analyses of principal coordinates (PCoA) performed on vOTU Bray-Curtis dissimilarities calculated across all viromes from the O horizon (A) and A horizon (B), with colors indicating treatment (field, control, or heat treatment) and shapes indicating virome type. UpSet plots of DNase-treated viromes in O horizon (C) and A horizon (D) and of nonDNase-treated viromes in O horizon (E) and A horizon (F), with filled circles indicating vOTU detection in at least two replicates of a particular treatment (field, control, or treatment temperature) and intersection size indicating the number of vOTUs with a given detection pattern. (G) DNA yields for DNase-(left facet) and nonDNase (right facet)-treated viromes from each horizon for each treatment condition. Box boundaries correspond to 25th and 75th percentiles, and whiskers extend to ±1.5x the interquartile range.

When considering the non-DNase viromes, recalling that these viromes likely contained genomes from compromised virions, the same trend was apparent in the O and A horizon, with most (43% for O and 37% for A) unique vOTUs shared across the ambient community and all treatment groups (in this case including 90°C). The ambient community and 60°C had the second most shared vOTUs (35% for O and 16% for A). The ambient community alone had the third most shared vOTUs in the O horizon (11%), but in the A horizon, the number of vOTUs shared among ambient samples (5.4% of total number of vOTUs) was not the third most-ranking intersection (Figures 3E&F). Consistent with increasing decay of viral particles in the 60°C and 90°C treatments from both horizons, there were significant differences based on temperature treatment on DNase-treated and non-DNase-treated viromic DNA yields (P < 0.05 by Kruskal-Wallis rank sum test) (Figure 3G). Together, these results suggest that treatment up to 30°C had no significant effect on viral particle abundance or viral community composition, but both DNA yields and community composition changed significantly at 60°C and again at 90°C. Differences between the ambient viral communities and the two higher temperature treatment groups were largely due to differences in the relative abundances of vOTUs present across treatments, rather than successional community changes, though there were a small number of vOTUs distinct to specific treatment conditions.

### Significantly differentially abundant vOTUs clustered into heat-sensitive and heat-tolerant trait groups

We performed differential abundance analysis on vOTUs in DNase-treated viromes (from likely intact virions) to determine which vOTUs were enriched or depleted as temperature increased. We identified 2,413 vOTUs (13% of total number of vOTUs in DNase-treated viromes) that were significantly differentially abundant across all pairwise temperature comparisons. We then tracked the change in the mean relative abundance of these vOTUs in all replicates within a temperature group and within a horizon over the temperature gradient. two distinct trend groups emerged for both horizons: vOTUs that increased in abundance as temperature increased, labeled as “heat tolerant,” and vOTUs that decreased in abundance as temperature increased, labeled as “heat sensitive” (Figures 4A&B). We then investigated the extent to which membership in each trend group was preserved between the two soil horizons. While the majority of vOTUs (74%) were uniquely found in a particular trend group in only one of the soil horizons, 454 vOTUs were heat-sensitive (19% of total) and 149 heat-tolerant (6% of total) in both the O and A horizons (Figure 4C).

**Figure 4.**
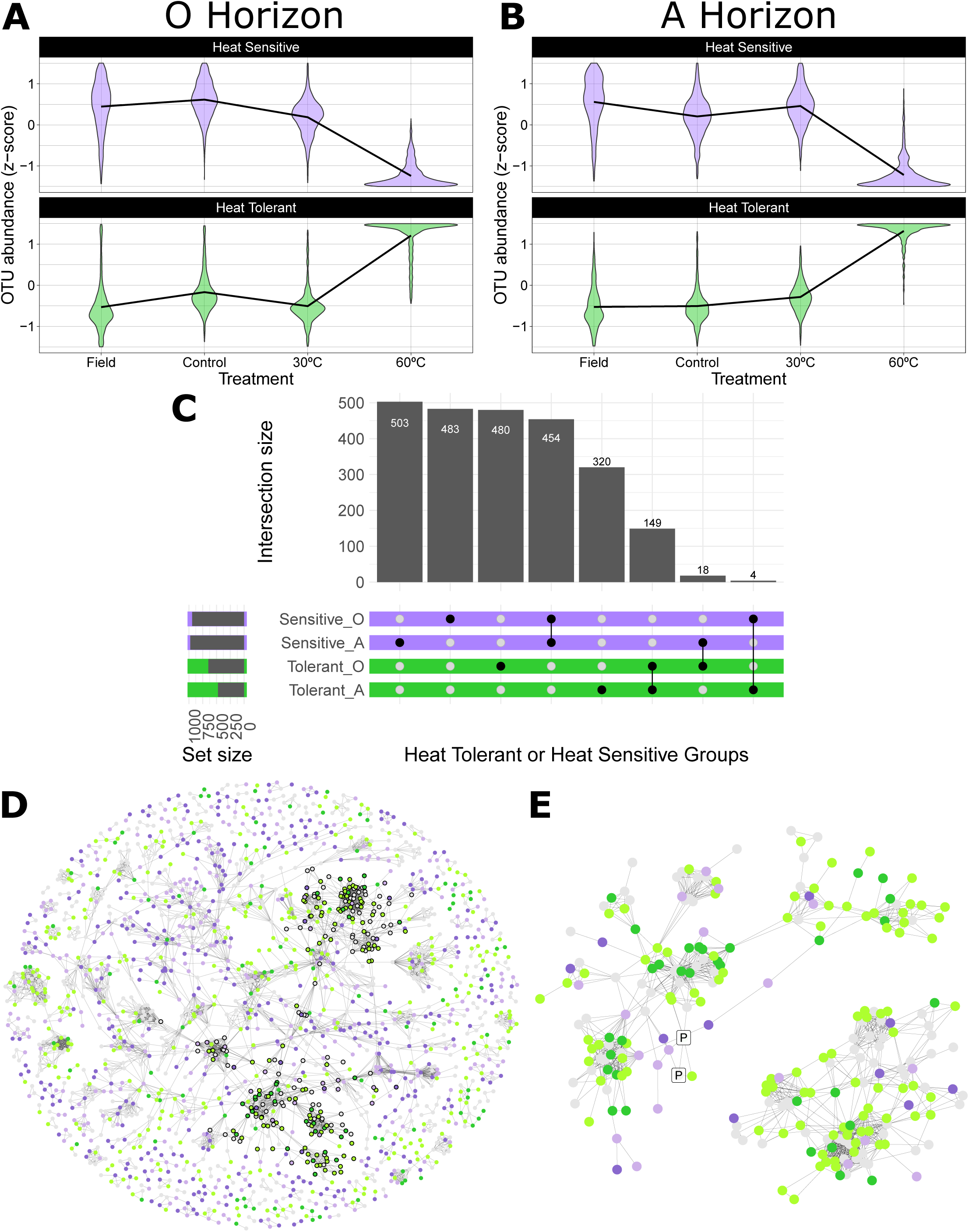
Patterns of differentially abundant vOTUs based on heat treatments. Violin plot visualizing the distribution of vOTUs (in terms of relative abundance, represented as a z-score) detected as significantly differentially abundant based on heat treatment in DNase-treated viromes from O horizon samples (A) and A horizon samples (B), with vOTUs that were enriched in the ambient samples (Field, Control, and 30°C) in the upper facet (“Heat Sensitive” trend group) and vOTUs that were enriched in the 60°C samples (“Heat Tolerant” trend group) in the lower facet. Trend line indicates mean z-score. (C) UpSet plot of differentially abundant vOTUs in DNase-treated viromes, with filled circles indicating membership in a trend group in a particular horizon and intersection size indicating the number of vOTUs with a given detection pattern. (D) Gene-sharing network displaying significant overlaps in predicted protein content (edges) between vOTUs (nodes). Node color represents membership in trend group of both (darker shade) or either (lighter shade) soil horizon, with absence of color indicating vOTUs that were not significantly differentially abundant based on temperature treatment. Black outlines around nodes indicate a subnetwork of local neighborhoods with significant overrepresentation of vOTUs in the “Heat Tolerant” trend group. (E) Gene-sharing subnetwork indicated in D, with nodes surrounded by a square indicating vOTUs that had significant overlap in their predicted protein contents with any RefSeq virus genome, with only two vOTUs that were significantly linked to RefSeq viruses, with both RefSeq viruses isolated from Proteobacteria hosts (tagged with letter “P”).

Finally, we investigated whether the trait of heat sensitivity or tolerance was taxonomically conserved by performing network analyses to place vOTUs into viral network neighborhoods. We identified 1,995 network neighborhoods, which consisted of vOTUs with at least one shared predicted protein with at least one other vOTU in the same neighborhood (Figure 4D). We identified 41 neighborhoods that were enriched in heat-sensitive vOTUs in both horizons, 31 enriched in heat-tolerant vOTUs in both horizons, 48 enriched in heat-sensitive vOTUs in only the A horizon, and 41 enriched in heat-tolerant vOTUs in only the O horizon. We did not identify any neighborhoods that were enriched in heat-sensitive vOTUs in only the O horizon, heat-tolerant vOTUs in only the A horizon, or containing vOTUs with mixed traits between the two horizons. To further investigate taxonomic associations of heat-tolerant vOTUs, we identified a subnetwork with clusters that had a significant proportion of these members. These clusters had a total of 250 nodes (vOTUs, Figure 4E), suggesting some taxonomic similarity among vOTUs that persist at higher temperatures. Only two RefSeq viruses were found in this subnetwork, and they were both viruses of Proteobacteria, suggesting that other viruses in the subnetwork might also infect Proteobacteria.

### Prokaryotic community responses to heat treatments were similar to those of viral communities

We assessed prokaryotic community responses to heat treatments by leveraging 16S rRNA gene amplicon data from both total DNA and non-DNase-treated viromes. Although the recovered prokaryotic communities were significantly different in the total and <0.22 µm (viromic) size fractions, the patterns in response to temperature were similar for each fraction in each horizon (Figures 5A&B). Like the viral communities, soil prokaryotic communities were most significantly distinct by soil source (soil horizon) (P < 0.001 by PERMANOVA) (Supplementary Figure 2A). Within each soil type, prokaryotic community composition also differed significantly with temperature, separating into the same three temperature groups as for the viral communities: field/control/30°C, 60°C, and 90°C. Similar to the viral communities, we can consider the prokaryotic ambient community to be best represented in the Field, Control, and 30°C samples, with significant compositional shifts as the temperature increased to 60°C and then 90°C (P < 0.001 for O horizon, for both sample types independently and P < 0.001 for A horizon for both sample types independently by PERMANOVA) (Supplementary Figures 2B-E).

**Figure 5.**
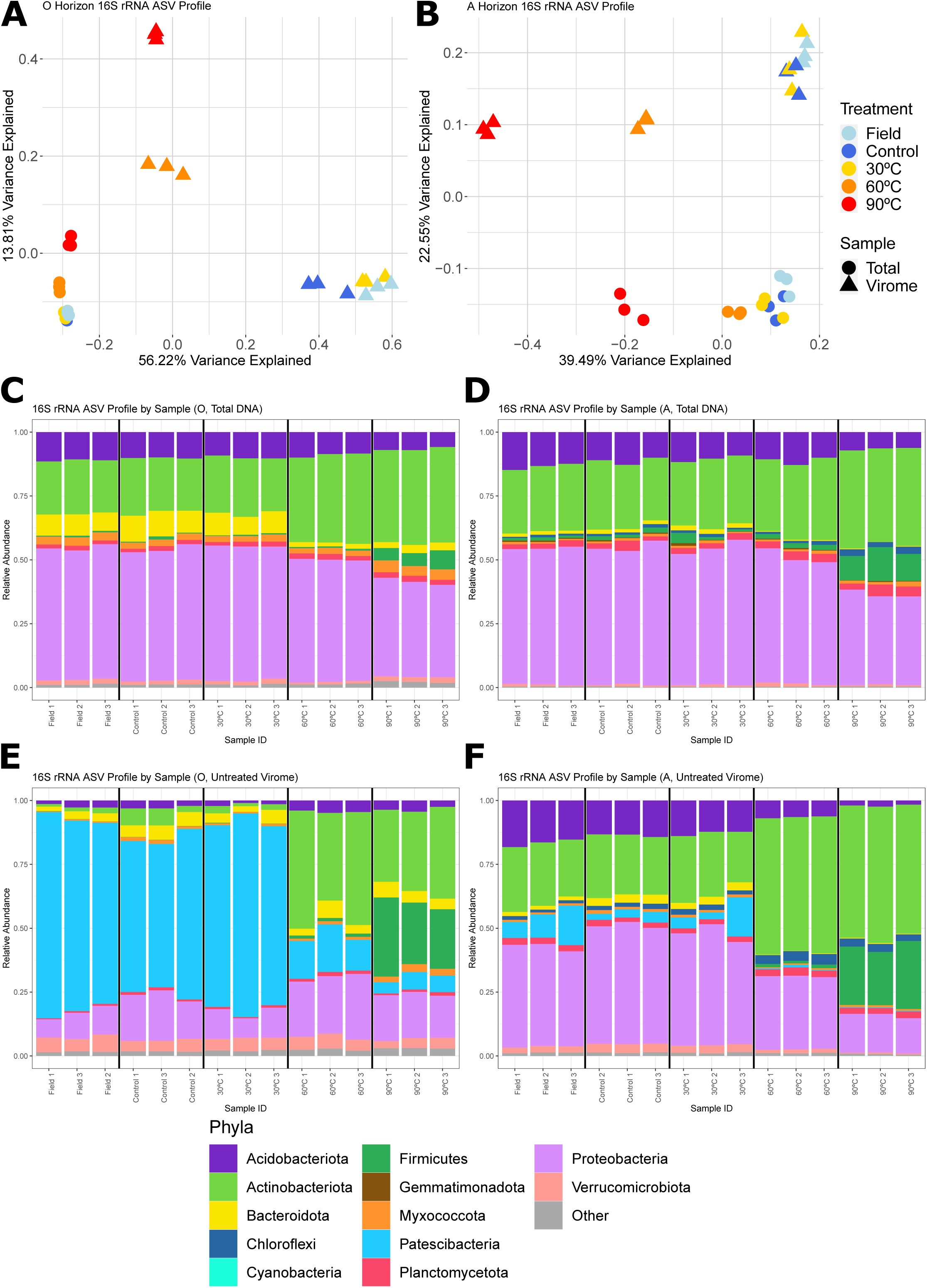
Prokaryotic community compositional trends according to heat treatment. Unconstrained analyses of principal coordinates (PCoA) performed on Bray-Curtis dissimilarities calculated from 16S rRNA gene Amplicon Sequence Variant (ASV) relative abundances in total DNA and nonDNase-treated virome sample types (shapes), with heat treatment indicated by color in the O horizon (A) and the A horizon (B). Phylum-level relative abundances in 16S rRNA gene profiles from total DNA amplicon libraries in the O horizon (C) and A horizon (D) and from nonDNase virome libraries in the O horizon (E) and horizon (F). Each stacked bar plot corresponds to a replicate within a treatment type, and the 12 most abundant phyla are colored, with all other low abundance phyla collapsed into an “other” group.

Underlying the prokaryotic community compositional patterns were taxonomic shifts that largely reflected similarity across the ambient conditions and changes in the relative abundances of specific taxa at 60°C and again at 90°C. In total DNA from the O horizon, the relative abundances of Acidobacteriota, Actinobacteria, Bacteriodota, Myxococcota, and Proteobacteria were significantly impacted by heat treatment (Kruskal-Wallis rank sum test, p < 0.05) (Figure 5C), as were all of those taxa, except for Myxococcota, in the A horizon (Kruskal-Wallis rank sum test, p < 0.05) (Figure 5D). For the >0.22 µm ‘viromic’ fraction in the O horizon, the relative abundances of the same taxa as in total DNA from the O horizon (Acidobacteriota, Actinobacteria, Myxococcota, and Proteobacteria), as well as Firmicutes and Patescibacteria, were significantly impacted by temperature treatment (Kruskal-Wallis rank sum test, p < 0.05) (Figure 5E). Finally, in the A horizon, >0.22 µm ‘viromic’ fraction, the relative abundances of 11 of the 12 most abundant phyla in the dataset (all but Planctomycetota) differed significantly by temperature treatment (Kruskal-Wallis rank sum test, p < 0.05) (Figure 5F). There were some consistent shifts across both soil horizons and sample type (total DNA or non-DNase treated virome), including an increase in the relative abundances of Actinobacteria and Firmicutes with increasing temperature. Additionally, Proteobacteria decreased in relative abundance as temperature increased, except for in the >0.22 µm ‘viromic’ fraction from the O horizon, where there was no significant trend. There were also distinct differences in each subgroup of soil horizon and DNA fraction. In the total DNA samples from the O horizon, there was a slightly higher enrichment of Bacteroidota in the ambient temperature group (on average 8.5% relative abundance), which decreased at higher temperatures (on average 1.8% at 60°C and 2.9% at 90°C). In the >0.22 µm ‘viromic’ fraction, the most notable taxon was the Patescibacteria (synonymous with Candidate Phyla Radiation), which are small-celled bacteria known to be enriched in 0.2 µm filtered soil samples (Nicolas et al., 2021; Santos-Medellín et al., 2023). The Patescibacteria dominated the prokaryotic sequences recovered from the ambient temperature group in the O horizon (on average 70% relative abundance) and were also detected at lower but still high abundance in ambient samples from the A horizon (7%), decreasing significantly in the O horizon (15% at 60°C and 5.8% at 90°C) and in the A horizon (0.6% at 60°C and 0.3% at 90°C) as temperature increased. Overall, prokaryotic communities exhibited similar beta-diversity trends to viruses, in terms of both differences by soil type (O vs. A horizon) and across the temperature gradient, suggesting that, in addition to the direct impacts of temperature, viral communities could be shifting in response to host communities on the timescale of this experiment.

To determine whether there was differential survival between viruses and their hosts, we first sought to link the two using iPHoP, a machine learning tool that predicts hosts for viruses of prokaryotes via a series of sequencing-based markers (Roux et al., 2023). Phylum-level taxonomic information of predicted hosts was assigned to 1,934 vOTUs, or 10% of the vOTUs in this dataset. Based on the Kruskal-Wallis rank sum test, there were significant differences across the temperature gradient in the relative abundance of vOTUs with a predicted host of Actinobacteria (p = 0.019) or Cyanobacteria (p = 0.02) in the O horizon and Actinobacteria (p = 0.02) or Proteobacteria (p = 0.02) in the A horizon (Supplementary Figures 1E&F). While we had intended to leverage the 16S rRNA gene sequencing data to compare specific, linked virus-host responses to temperature, meaningful data analysis was not possible, due to limited host prediction. We did find that most of the predicted host phyla were also found in the ASV profiles in the respective horizon.

## DISCUSSION

This study revealed a likely upper limit to viral persistence between 60 and 90°C for most, if not all, viral species in the studied soils. Given that DNA yields for DNase-treated viromes were below detection limits for all six samples that underwent heating treatment at 90°C and that these DNase-treated viromes likely captured predominantly intact virions (Santos-Medellín et al., 2023; Sorensen et al., 2021), we can infer substantial or complete depletion of intact soil viral particles at 90°C. The depletion threshold presumably occurred between 60°C and 90°C, or possibly with an upper limit as low as 83°C, which was the lowest maximum temperature achieved in the center of the microcosms incubated at 90°C. We can thus assume a maximum viability threshold between 83°C and 90°C for most soil viruses from these forest soils. This threshold is aligned with a study that focused on 24 viruses of thermophilic *Bacillus* strains isolated from compost, soil, silage, and rotting straw, in which the ability to form plaques was possible at 50°C but not at 70°C (Sharp et al., 1986). Additionally, while studies have shown heat resistance in viral isolates at above 90°C (Guglielmotti et al., 2012), there is evidence that virus inactivation occurs faster in complex matrices, such as soil (Bertrand et al., 2012). Performing heating treatments with higher resolution between 60°C and 90°C and at multiple heating durations will likely reveal more precise temperature thresholds for different groups of dsDNA prokaryotic viruses in soil.

Although data are limited for viruses, thermotolerance for prokaryotes is much better understood, and in general, the inferred viral thermotolerances here are consistent with those of prokaryotes. In response to heat treatment, within both soil types, both viruses and prokaryotes clustered into three groups, with community composition similar across the ambient community (field, control, and 30°C samples) and at 60°C and again at 90°C. Changes in vOTU relative abundances better explained separation among viral communities along the temperature gradient than did compositional turnover, with most viral species detectable across the temperature gradient. This suggests that thermotolerance up to at least 60°C is widespread among most viral ‘species’ in these soils. Since the largest intersection of vOTUs was present in all treatment groups in the DNase-treated viromes (which included all but the 90°C samples), we infer that most vOTUs in these soil samples were tolerant at 60°C. The idea of the ambient viral and prokaryotic communities having high thermotolerance seems to be supported by other studies focused on microbial populations. For example, in composting systems where the optimal growth temperatures for most strains were found to be 30 to 40°C, thermotolerance (up to 60°C, the maximum study temperature) was ubiquitous among all major phyla (Actinobacteria, Firmicutes, Proteobacteria), with most microorganisms considered eurythermal (adaptable to and able to tolerate a wide range of temperatures) (Moreno et al., 2021). In terms of prokaryotic trends in response to temperature change, the increase in relative abundance of Firmicutes at 90°C is also consistent with prior studies, with members of the Firmicutes exhibiting persistence in hostile habitats, spore-forming capabilities, and a relatively high growth rate, comprising the majority of thermotolerant strains with high thermal plasticity (Moreno et al., 2021). The increase in relative abundance of Chloroflexi at 90°C in this study is consistent with increasing relative abundance of this phylum with increasing temperature in Costa Rican hot springs that reached up to 63°C (Uribe-Lorío et al., 2019). The significant decrease of Patescibacteria at higher temperatures (a result only evident in the prokaryotic data from the viromes, which enrich for small cells) also may be due to the lower number of functional genes related to stress response in their small genomes, as compared to other bacteria with larger genome sizes (Tian et al., 2020). The detection of viruses in this study that were more significantly abundant at 60°C suggests that some of these viruses may exhibit heat tolerance, similar to prior discoveries of heat-tolerant prokaryotic phyla. However, the connection between viruses and their hosts and whether they exhibit differential or parallel survival with increasing temperature requires further investigation.

In this study, we found that both viral and prokaryotic communities exhibited similar beta-diversity patterns according to soil source. Community composition differed most significantly by soil source (O or A horizon). Viral community differences between O and A horizons have not been extensively studied in other soils, but viral communities are known to differ by soil depth (Emerson et al., 2018; Liang et al., 2019; Ter Horst et al., 2021), consistent with differences between horizons found in this study. For prokaryotes, the observed compositional differences were consistent with known differences between soil horizons. For example, as in this study, Acidobacteria, Proteobacteria, and Actinobacteria are ubiquitous in forest organic and mineral soils, with Bacteroidetes more prominent in organic soils and Firmicutes and Chloroflexi more prominent in mineral soils (Baldrian et al., 2012; Uroz et al., 2013). While considered rare in soil (Liu et al., 2020), here Patescibacteria represented a substantial proportion (up to 80%) of the prokaryotes detected in the O horizon ‘viromic’ (< 0.22 µm) fraction. They were also more prominent in organic compared to mineral soils, which may be due to their tendency to metabolize simple metabolites like glucose and pyruvate which may be more available in organic matter rich soils (Tian et al., 2020).

## DATA AVAILABILITY

All raw sequences have been deposited in the NCBI Sequence Read Archive under the BioProject accession <u>PRJNA1093237</u>. The database of dereplicated vOTUs is available at https://zenodo.org/records/11043663. All scripts are available at https://github.com/seugeo/soilheating.

## ACKNOWLEDGEMENTS

We thank Brandon Matsumoto for help with sample collection, staff at the Blodgett Forest Research Station (Kane Russell, Amy Mason, and Ariel Thomson) for advising on field sampling logistics and coordinating site visits, and resource support from the Traxler Lab at UC Berkeley (Monika Fischer, Neem Patel). Thanks to Valerie Eviner and Jorge Rodrigues for helpful comments on a draft version of the manuscript. Funding for this work was provided by the U.S. Department of Energy (DOE), Office of Science, Office of Biological and Environmental Research (BER), Genomic Science Program, award number DE-SC0021198 (grant to JBE). SEG was also supported by the National Science Foundation Graduate Research Fellowship.

