## Supplementary Figures for "Virome responses to heating of a forest soil suggest that most dsDNA viral particles do not persist at 90°C"

**Supplementary Figure 1. Viral community composition and predicted host abundances based on heat treatments** Unconstrained analyses of principal coordinates (PCoA) performed on vOTU Bray-Curtis dissimilarities calculated across DNase-treated viromes from the O horizon (**A**) and A horizon (**B**) and across non DNase-treated viromes from the O horizon (**C**) and A horizon (**D**), with colors indicating treatment (field, control, or heat treatment) and shapes indicating virome type. Relative abundances of vOTUs with a phylum-level predicted host in DNase-treated viromes in the O horizon (**E**) and A horizon (**F**). Each stacked bar plot corresponds to a replicate within a treatment type, and the 5 most abundant host phyla are colored, with all other low abundance host phyla collapsed into an “other” group.

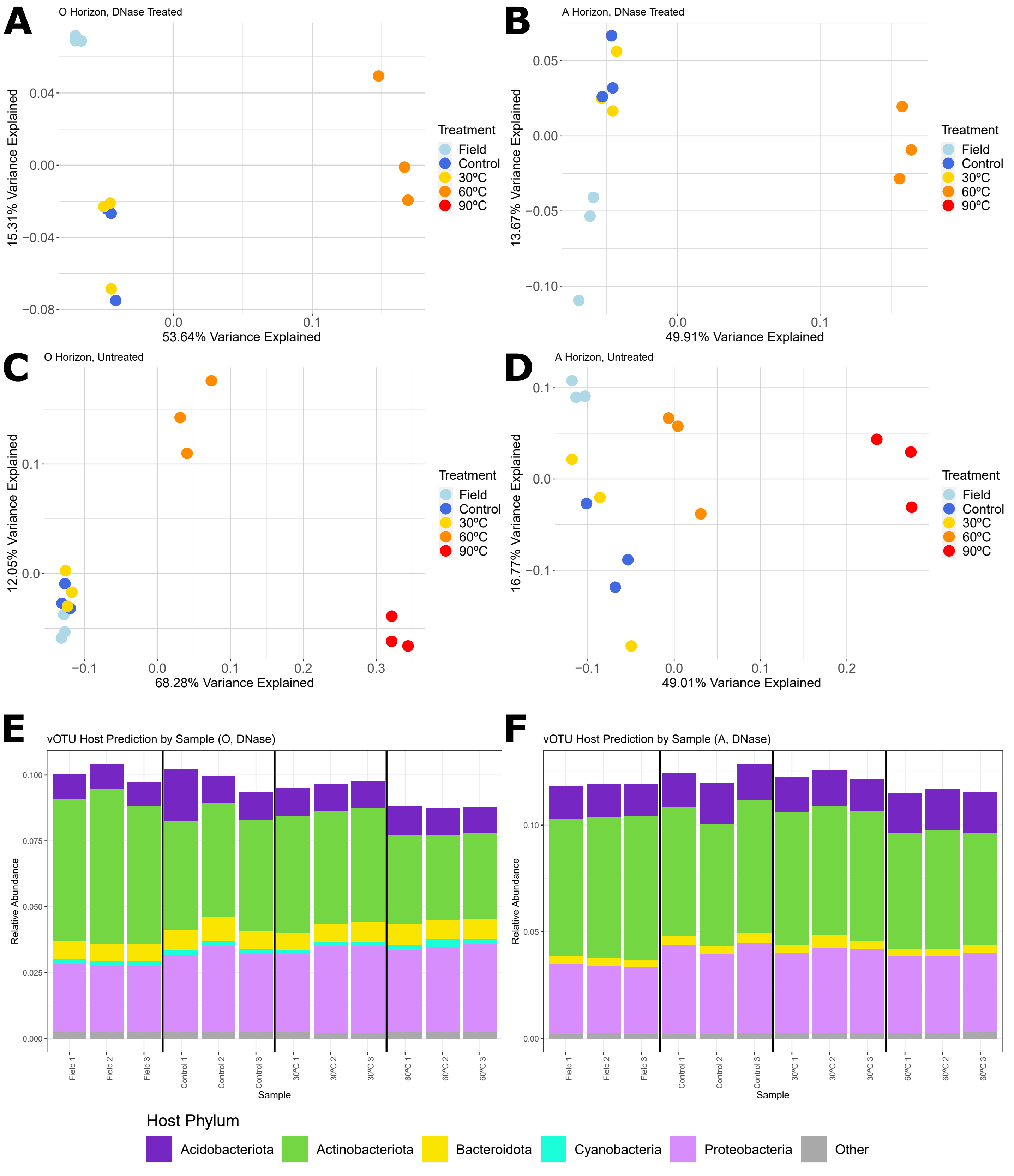

**Supplementary Figure 2. 16S rRNA gene ASV community composition based on soil type and heat treatments** Unconstrained analyses of principal coordinates (PCoA) performed on vOTU Bray-Curtis dissimilarities calculated across all total DNA and non-DNase-treated viromes, with colors indicating horizon and shapes indicating sample type (**A**). PCoA performed on vOTU Bray-Curtis dissimilarities calculated across total DNA samples in the O horizon (**B**) and A horizon (**C**) and across non-DNase-treated viromes in the O horizon (**D**) and A horizon (**E**), with color indicating treatment (field, control, or heat treatment) and shape indicating sample type.

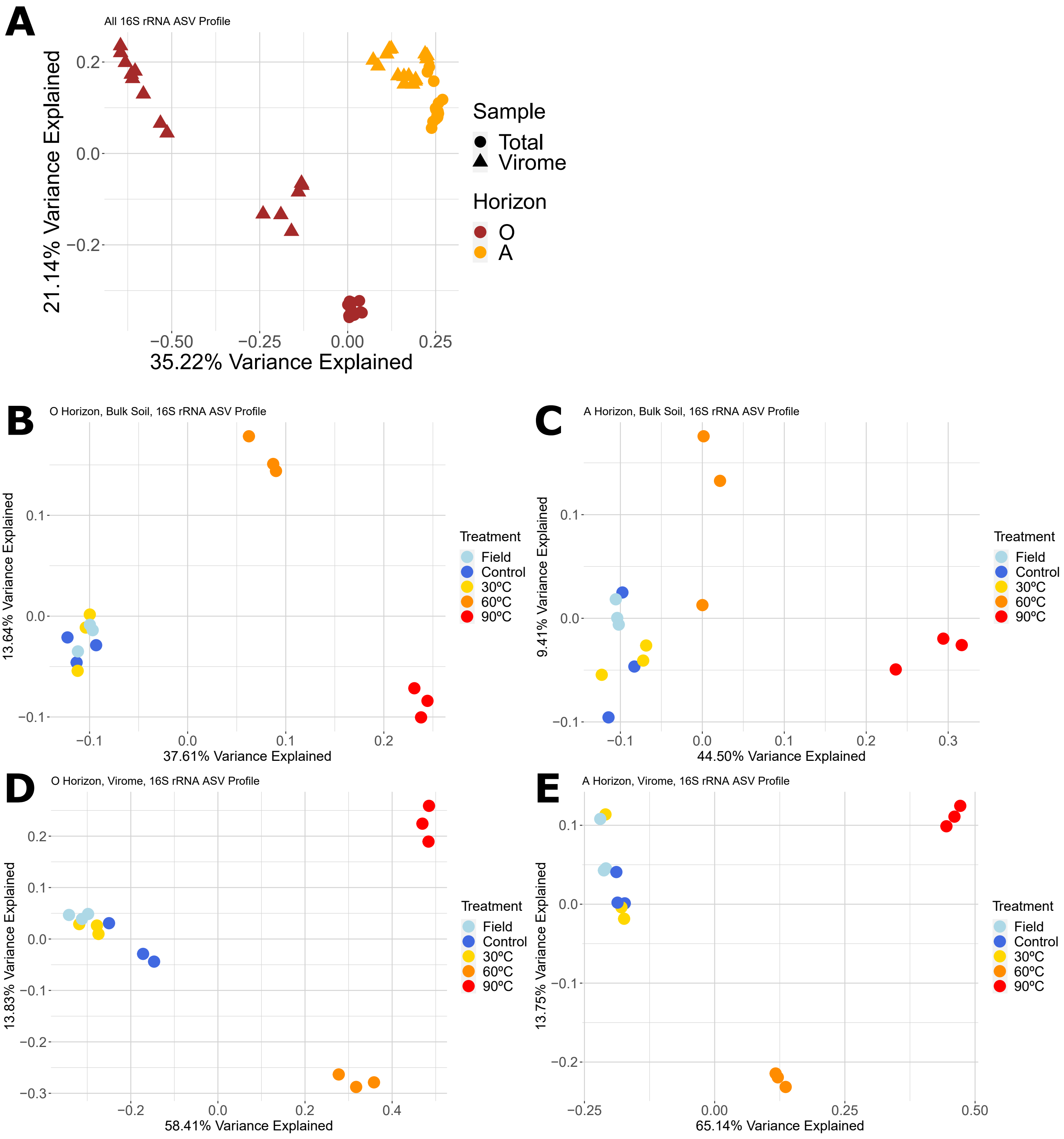
